# Implementation of a multiplex PCR amplfication system combined with next-generation genome sequencing to decipher the circulation of Human coronavirus 229E lineages in Southern France

**DOI:** 10.1101/2025.03.12.639729

**Authors:** Houmadi Hikmat, Justine Py, Céline Boschi, Emilie Burel, Lorlane Le Targa, Matthieu Million, Lucile Lesage, Aurélie Morand, Bernard La Scola, Philippe Colson

## Abstract

Coronaviruses are known to evolve rapidly and be prone to new virus emergence. Human coronavirus-229E is one of the seven coronaviruses currently known to cause respiratory symptoms in humans. Genomic data are very scarce for this virus.

Here, we implemented an in-house multiplex PCR strategy to amplify HCoV-229E genomes from nasopharyngeal samples diagnosed as positive for this virus RNA in Southeastern France. Then, next-generation sequencing was performed using Nanopore or Illumina technologies on Gridion or Novaseq 6000 instruments, respectively. HCoV-229E genomes were assembled and analyzed using MAFFT, MEGA, Itol, Nexstrain and Nextclade softwares.

Thirty-one PCR primer pairs were designed to amplify overlapping fragments of the HCoV-229E genome. They allowed obtaining 123 genomes, which were classified in an emerging HCoV-229E lineage first reported in China, with two sublineages being delineated. Relatively to genome NC_002645.1 dating back to 1962, regarding nucleotide mutations, 1,167 substitutions, 72 insertions and 34 deletions were detected in viral genomes obatined here, while regarding amino acid mutations, 415 susbstitutions, 39 deletions and 14 amino acid insertions were detected. The genes with the greatest diversity were S (that encodes the spike protein) then Nsp3. Signature mutations were identified for the two sublineages.

In summary, we almost doubled the set of HCoV-229E genomes available worldwide and provided the first genomes from France. Further studies are needed to strengthen the knowledge about the phylogenomics and evolutionary dynamics of this neglected respiratory virus, and about their specificities, which may also purvey clues to contribute improving knowledge for all other human coronaviruses.

## INTRODUCTION

Human coronavirus-229E (HCoV-229E) was the first discovered, in 1966, of the seven coronaviruses currently known to infect and cause respiratory diseases in humans^1,2^. Infections are most often associated with mild clinical symptoms but can be severe and life-threatening in children, elder people and in case of underlying illness^3^. They show a seasonality in temperate countries with the greatest incidence during winter and spring^4,5^.

HCoV-229E is classified in genus *Alphacoronavirus*^6^. Its genome is a single-stranded positive sense RNA with an approximate size of 27 kilobases (kb). Its first two-thirds encode non-structural proteins (namely, NSP1 to NSP16) and the remaining third encodes structural proteins including the spike (S), envelope (E), membrane (M) and nucleocapsid (N) proteins. An accessory protein is encoded by a gene located between those encoding the spike and the envelope ^7–9^. HCoV-229E is classified into six genotypes named 1 to 6, while an emerging lineage was reported in China in 2023^10,11^. Aminopeptidase-N is the primary cell surface receptor for this virus^12^. Natural and intermediate hosts for HCoV-229E are deemed to be bats and alpaca, respectively^1,13^.

The HCoV-229E genome was primarily studied betweeen 1978 and 2001^14–16^ and the first complete genome sequence from a clinical isolate was described in 2012^17^. Still, there is currently a huge discrepancy between the million SARS-CoV-2 genomes available in worldwide databases^18^ (https://gisaid.org/; https://www.ncbi.nlm.nih.gov/genbank/) and the only 130 genomes (as of 01/01/2024) available in the NCBI Genbank nucleotide sequence database. Besides, none of these genomes originated from France. There is therefore a considerable paucity of data and, consequently of understanding of HCoV-229E genetic diversity and evolution. Hence, here we aimed to sequence and analyze HCoV-229E genomes from respiratory samples that had been diagnosed as HCoV-229E RNA-positive in Marseille, Southeastern France.

## MATERIALS AND METHODS

### Respiratory samples

Next-generation sequencing (NGS) of HCoV-229E genomes was carried out retrospectively from remains of nasopharyngeal samples that had been sent to our clinical microbiology laboratory at University and Public Hospitals of Marseille, Southeastern France, for the purpose of routine diagnosis of respiratory infections in the setting of clinical routine management, and that were stored at −20°C or −80°C after processing. HCoV-229E RNA testing had been carried out by multiplex real-time reverse-transcription (RT)-PCR (qPCR), as previously described^19^.

### PCR primer design and PCR amplification of overlaping regions covering the whole genomes

All near complete or complete HCoV-229E genomes available from GenBank (https://www.ncbi.nlm.nih.gov/genbank/)^20^ as of 28/02/2022 were retrieved. Recovered genomes were aligned using the MAFFT software^21^. PCR primers targeting the most conserved regions of the genomes were then designed using the Gemi software^22^ to implement a PCR amplification primer set that enables generating overlaping amplicons covering the whole genome sequence, following the ‘ARTIC’ strategy used for instance for SARS-CoV-2 genomes (https://artic.network/ncov-2019/ncov2019-bioinformatics-sop.html). The list of PCR primers and the PCR conditions used for HCoV-229E genome amplification are provided in Table 1.

**Table 1.**
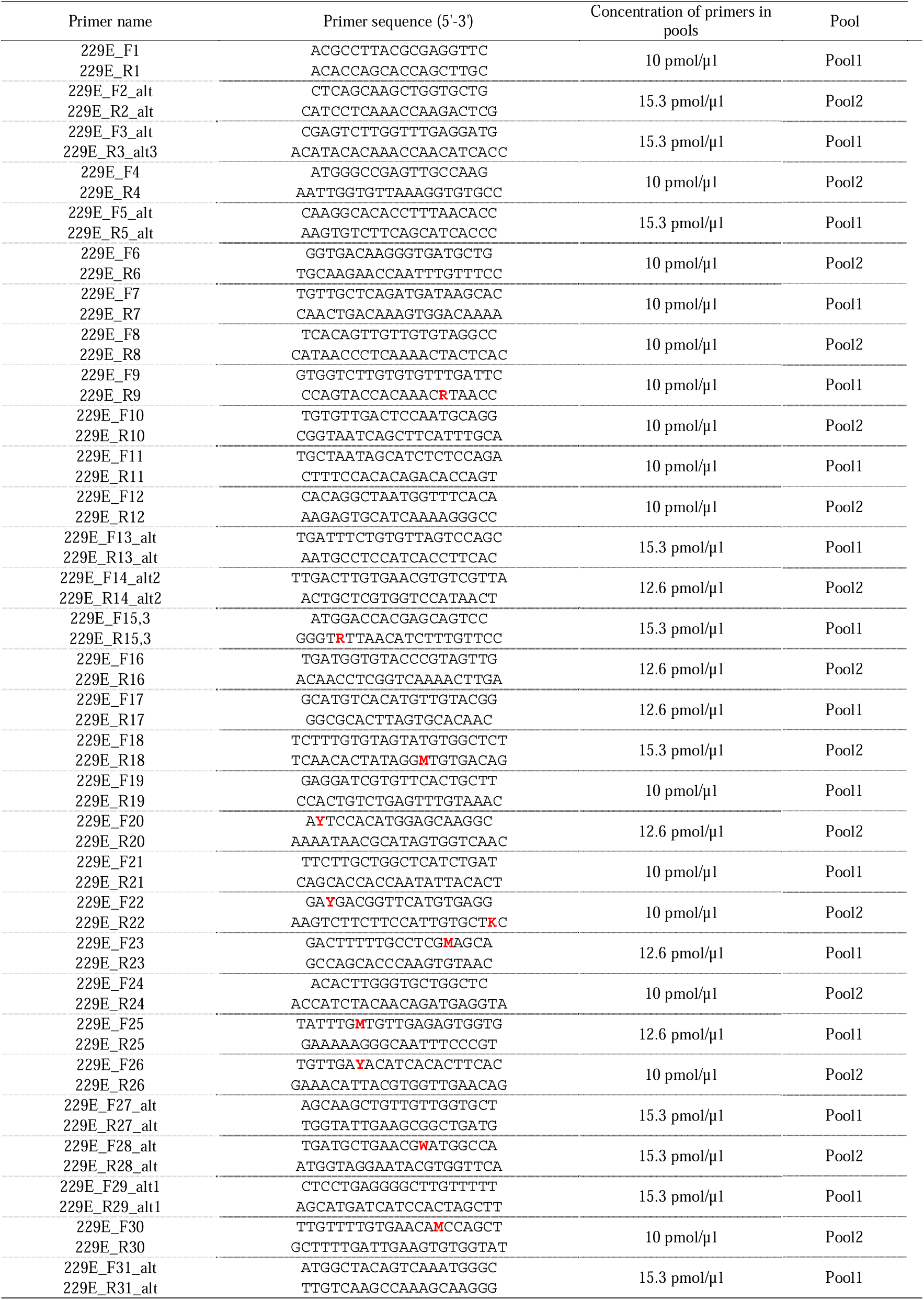
PCR primers and conditions used for the amplification of HCoV-229E genome fragments. RNA was extracted by the KingFisher Flex system (Thermo Fisher Scientific, Waltham, MA, USA) according to the manufacter’s protocol. Then RNA extracts were amplified by standard RT-PCR using the SuperScript III One-Step RT-PCR Kit with Platinum Taq High Fidelity (Invitrogen, Life Technologies, Carlsbad, CA, USA) and designed PCR primer pairs used in two pools not to have amplicon overlaps. PCR amplification included a reverse transcription step according to the following reaction: 12.5 µL of 2X mix, 0.7 µL of enzyme, 0.5 µL of each PCR pool, 3 µL of nucleic acid extract, and 8.3 µL of water. The PCR protocol included a preliminary reverse transcription step at 50°C for 25 minutes, followed by an initial denaturation at 95°C for 2 minutes, then by 39 PCR cycles including denaturation at 95°C for 15 sec, hybridization at 58°C for 45 sec, and elongation at 70°C for 1.5 min. Finally, a final elongation step at 70°C for 5 min was carried out. The PCR amplicons were purified on a NucleoFast 96-well plate (Macherey-Nagel, Hoerdt, France) with an elution with 40 µL of pure water. Post-PCR amplification, amplicons from the two primer pools were mixed. Degenerated nucleotides are indicated by a red font.

### Next-generation sequencing

For the step of testing designed PCR primers and PCR conditions, NGS used the Oxford Nanopore technology (ONT), with the Ligation sequencing kit SQK-LSK109 then library deposit on a SpotON flow cell Mk I, R9.4.1 and a GridION instrument, according to the manufacturer’s protocols (Oxford Nanopore Technologies Ltd., Oxford, UK). Thereafter, we performed NGS on RNA extracts obtained using the KingFisher Flex system (Thermo Fisher Scientific, Waltham, MA, USA) from available remains of nasopharyngeal samples that had been diagnosed as HCoV-OC43 RNA-positive in our laboratory. At this step, NGS was carried out using ONT, and Illumina technology on a NovaSeq 6000 instrument, with the CovidSeq protocol (Illumina Inc., San Diego, CA, USA) but with the replacement of the Covid-19 ARTIC PCR primers by the PCR primers we previously designed and according to the PCR conditions we previously set up. Loading procedure on a NovaSeq 6000 SP flow cell followed the NovaSeq-XP workflow and a procedure previously described^23^ with a reading of 2×50 nucleotides.

### Processing and bioinformatic analyses of NGS reads and viral genomes

Genome sequences were assembled by mapping on the HCoV-229E genome GenBank accession no. LC654445.1 (Fukushima_H829_2020 isolate) with Minimap2^24^. Samtools13 was used for soft clipping of Artic primers and removing sequence duplicates^25^. Consensus genomes were generated using Sam2consensus (https://github.com/edgardomortiz/sam2consensus). A phylogenetic tree was created with the MEGA^26^ software (v.11) using the Neighbor-Joining and Maximum composite likelihood parameter methods with 1,000 replicates after sequence alignment with the MAFFT software^21^. All HCoV-229E genomes available from GenBank including those corresponding to genogroups were incorporated in the phylogeny reconstruction. The Itol tool ^27^ was used to visualize the phylogenetic tree. Nextstrain ^28^ and Nextclade^29^ tools were adapted to enable identifying viral lineages and mutations. Nucleotide and amino acid diversity was obtained relatively to the HCoV-229E reference genome no. NC_002645.1.

## RESULTS

A total of 524 nasopharyngeal samples had been diagnosed as HCoV-229E RNA-positive in our institution between January 2017 and October 2022, but only remains for 195 of them, which had been collected between December 2022 and March 2022, were available as stored frozen and in sufficient volume (Figure 1). These 195 specimens were tested using a multiplex PCR amplification strategy to amplify the genome per short overlaping fragments for further NGS. After designing and testing individually possible PCR primer pairs, 31 primer pairs generating amplicons covering the entire HCoV-229E genome were selected (Table 1); primer concentration in PCR ranged from 10 to 15.3 µM. These PCR primer pairs were used in two separate pools to prevent hybridization of generated amplicons. They allowed obtaining 123 genomes with a completion corresponding to ≥80% coverage of reference genome NC_002645.1; mean (±standard deviation) coverage was 92.14%±0.05 (range, 80.0-98.0%). qPCR cycle threshold values (Ct) were available for 75 of the samples from which these 123 genomes were obtained. Mean Ct was 21.2±4.0 (15.0-29.0). Regarding the 72 other specimens for which HCoV-229E genomes could not be obtained with previously defined criteria, their mean Ct was 26.3±2.7 (22.0-30.0).

**Figure 1.**
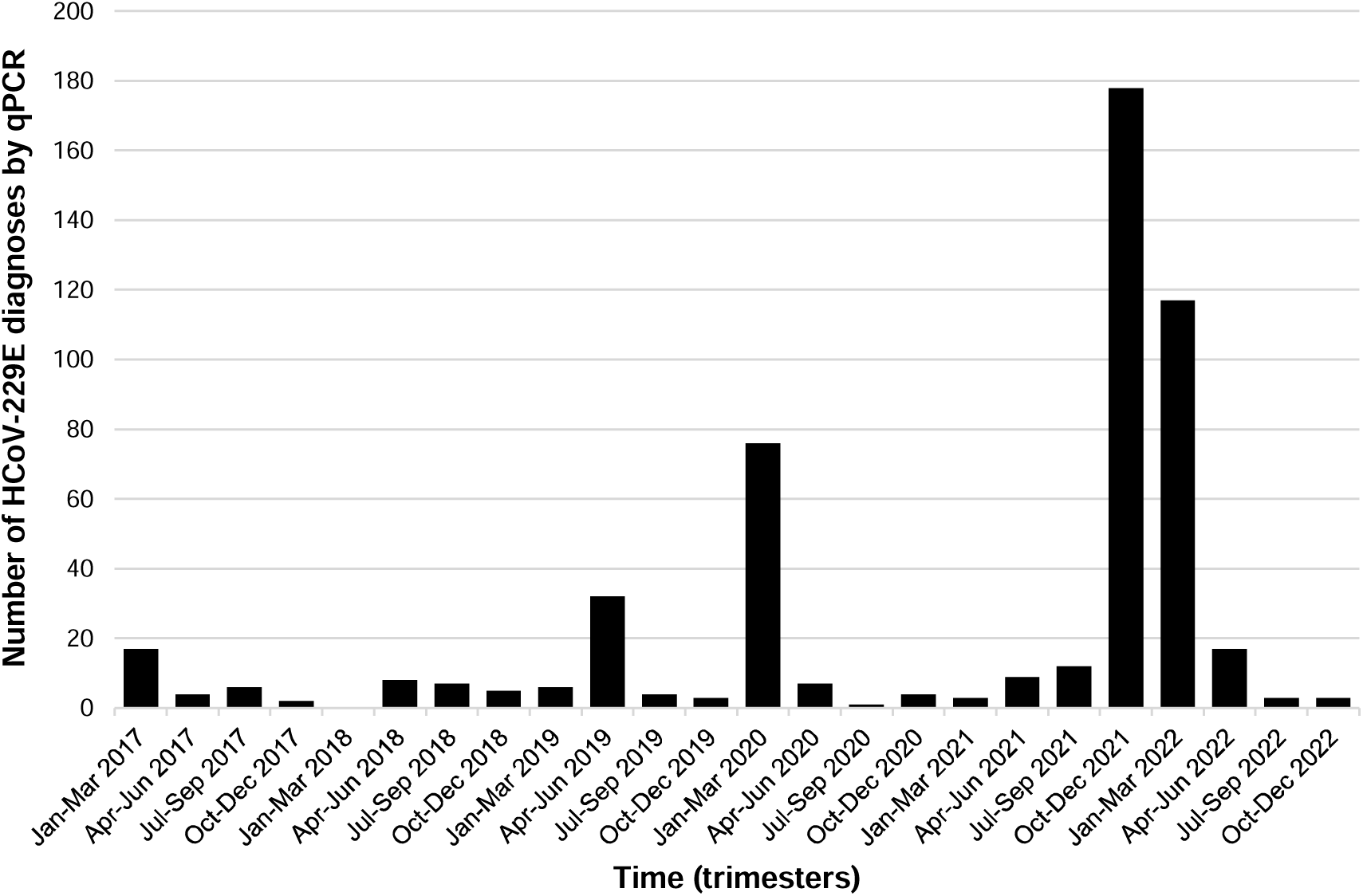
Temporal distribution of HCoV-229E RNA-positive nasopharyngeal samples that had been collected from patients between 2017 and 2022

The 123 near-full genomes were obtained from nasopharyngeal samples collected between March 2021 and March 2022. They were classified by the Nextclade tool and the phylogeny reconstruction as belonging to an emerging lineage reported in China in 2023^11^, which was designated as a putative genotype 7 in two recent reports^30,31^(Figure 2). Based on the phylogenetic analysis, this emerging lineage comprised two sublineages, one of which appeared to match with previously designated sublineages 7b^30,31^ whereas there seems to be discrepancies between matches for the second sublineages and previously designated submineage 7a^30,31^. Whatever, 30 of the 123 genomes obtained here belong to a sublineage “a” and 93 belong to a sublineage “b” (Figure 2), revealing that these two sublineages co-circulated in our geographical area, with a predominance of viruses of sublineage b (Figure 3). Sublineage a was detected since March 2021 while sublineage b was detected since September 2021. For the four months during which the number of genomes obtained from collected specimens were above 10, the proportion of genomes of sublineage a decreased from 32% in November 2021 to 7% in February 2022.

**Figure 2.**
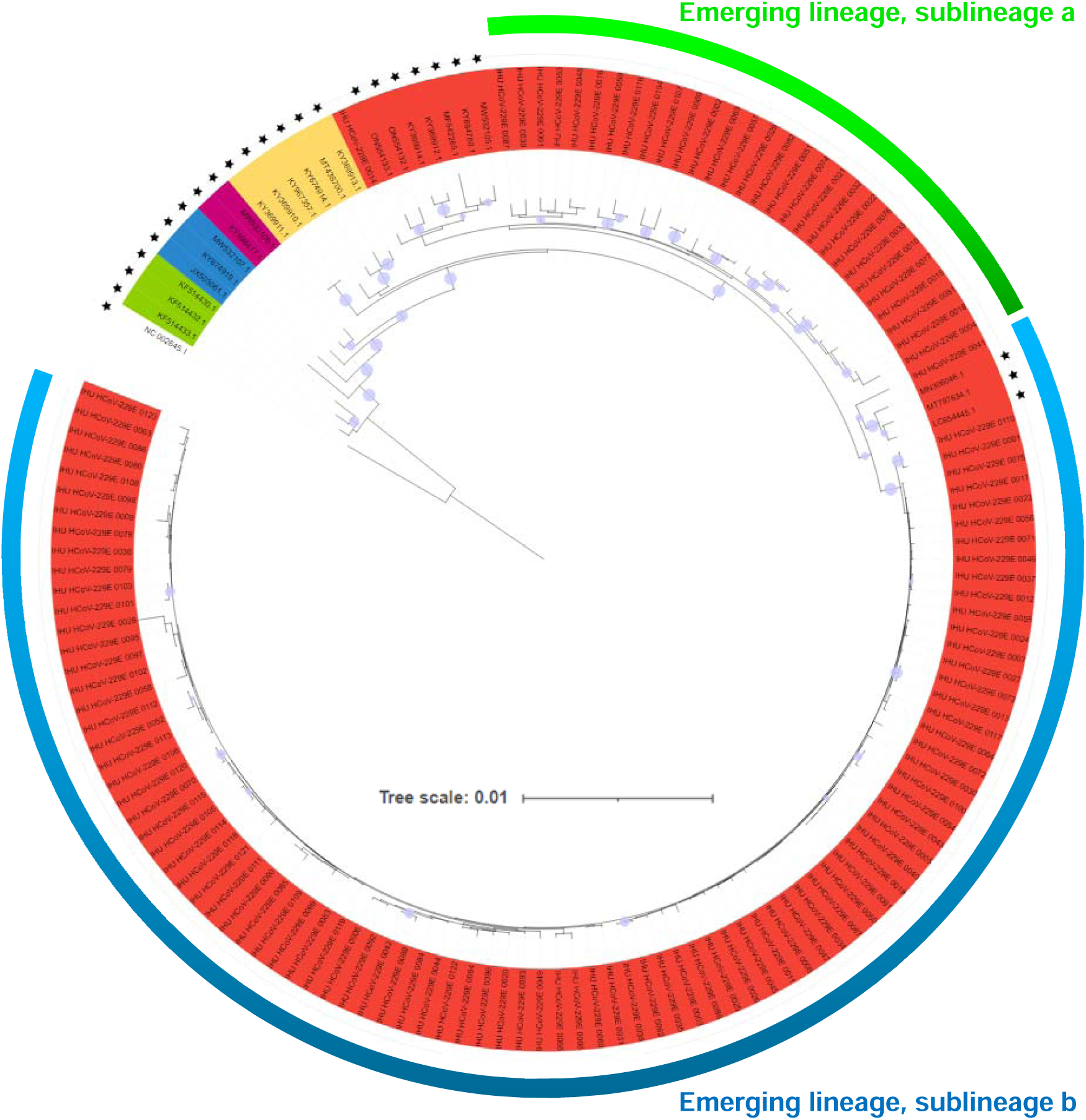
Phylogeny reconstruction based on HCoV-229E genomes recovered in the present study or available from GenBank, including genomes from the different previously delineated lineages or sublineages Genomes from GenBank are indicated by a black star.

**Figure 3.**
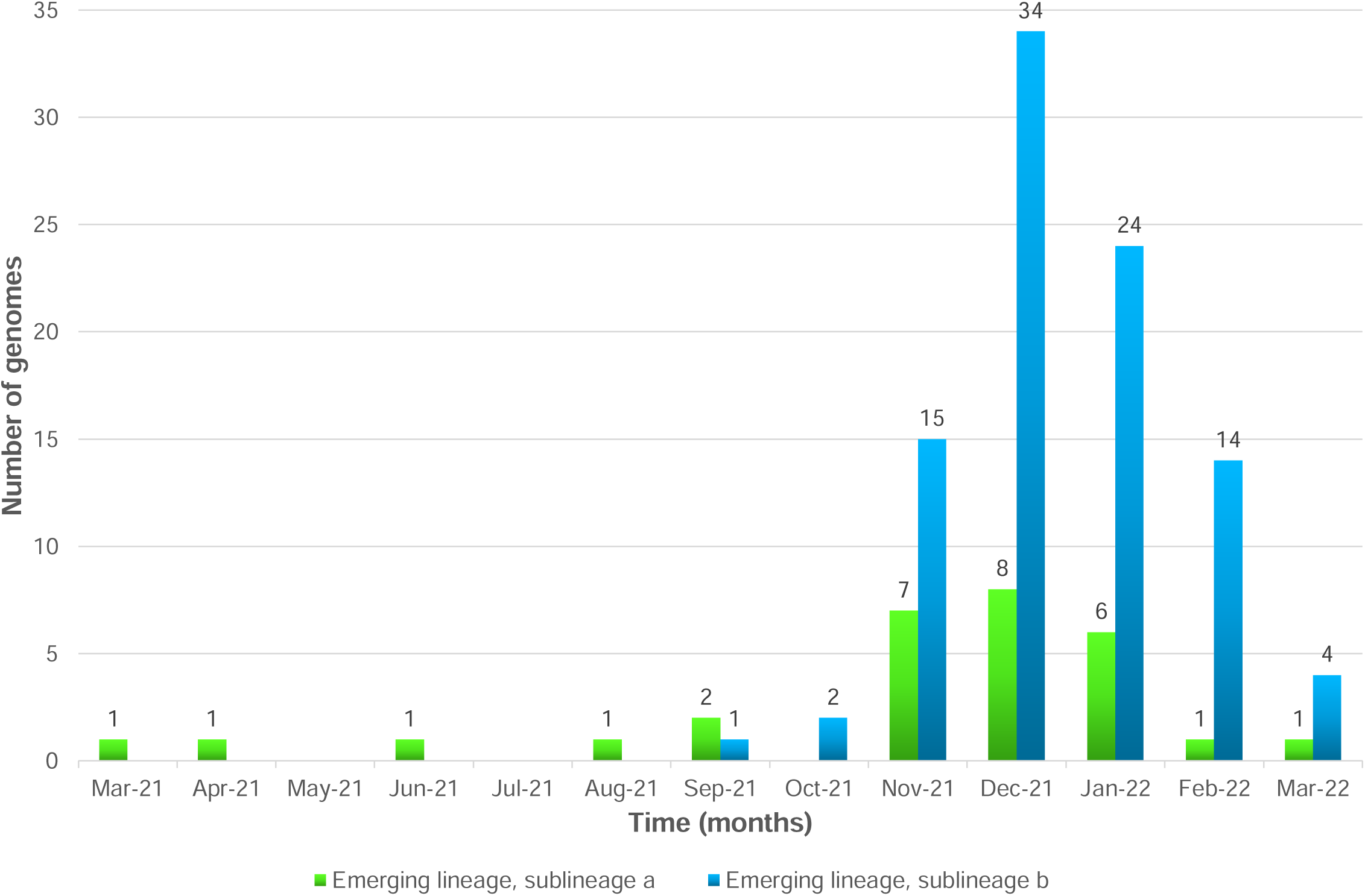
Temporal distribution of the two subgenotypes of HCoV-229E detected in the present study

The nucleotide and amino acid diversity of the 123 HCoV-229E genomes was determined relatively to genome NC_002645.1 first submitted to Genbank in 2001. Regarding nucleotide mutations, 1,167 substitutions, 72 insertions and 34 deletions were detected in these genomes. Regarding amino acid mutations, 415 substitutions, 39 deletions and 14 amino acid insertions were detected. Of the 415 amino acid substitutions, only 211 were present in ≥5 genomes. These 211 amino acid substitutions were found in several genes, including in the Nsp1, Nsp2, Nsp3, Nsp4, Nsp6, Nsp8, Nsp9, Nsp10, Nsp11, Nsp12, Nsp13, Nsp14, Nsp15, spike, ORF4a, E, M, and N genes. A total of 78 (37%) of these 211 amino acid substitutions were in the spike. They displayed various prevalence (Figure 4, Table 2). Seventy-one were already reported, being mentioned as new substitutions in 13 cases^11^. Spike deletions S:V353- and S:Y354- were observed in all genomes obtained here, while S:A352- was observed in all but four genomes (97%) that harbor four other deletions: S:A355-, S:N356-, S:V357- and S:G358- (Table 2). Two other deletions were present in NSP3 (NSP3:L105-, NSP3:P106-) of all genomes (except one for which these codons were not covered).

**Figure 4.**
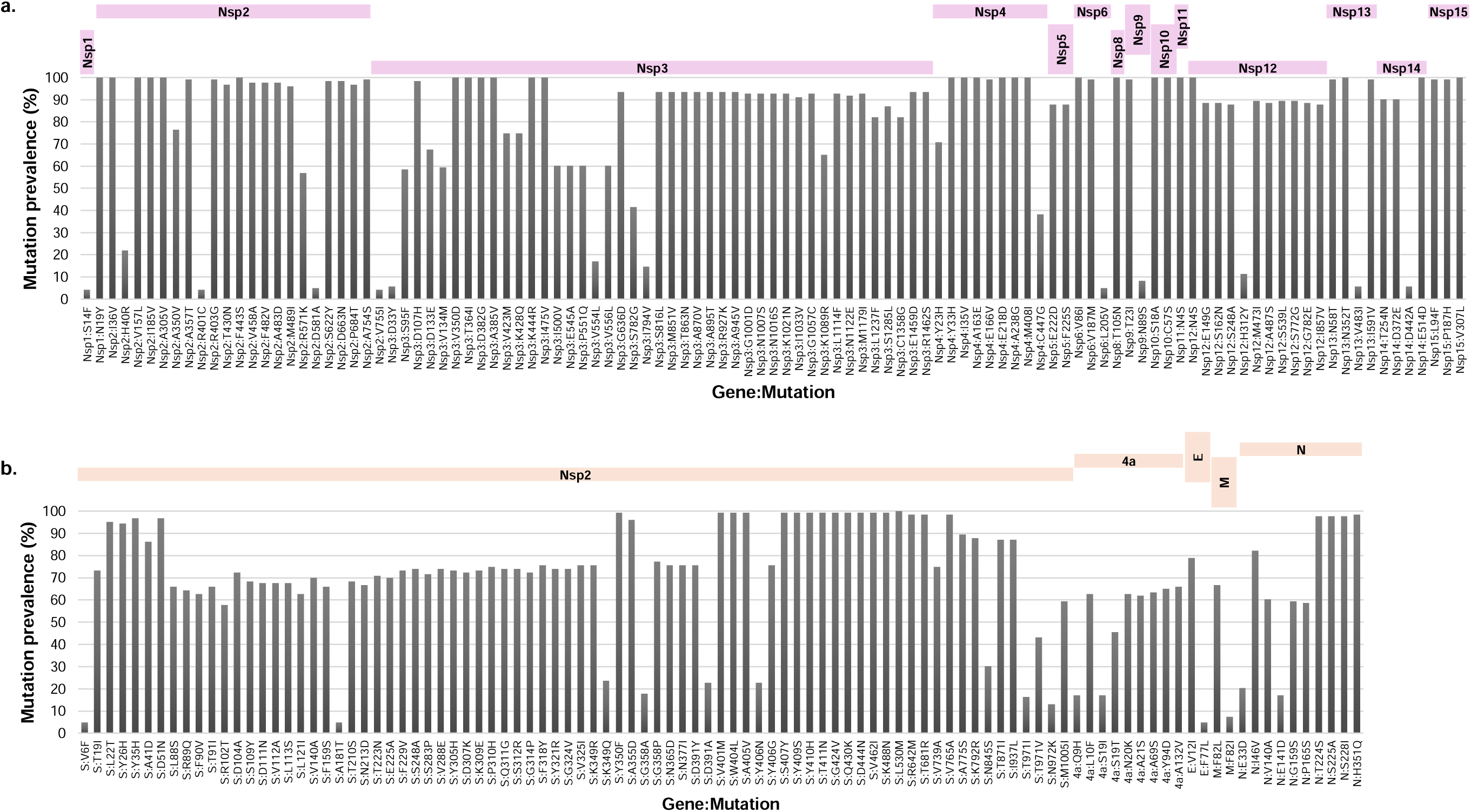
Amino acid substitution prevalence in non-structural (a) and structural (b) genes E, envelope; M, Membrane; N, Nucleocapsid; S, Spike.

**Table 2.**
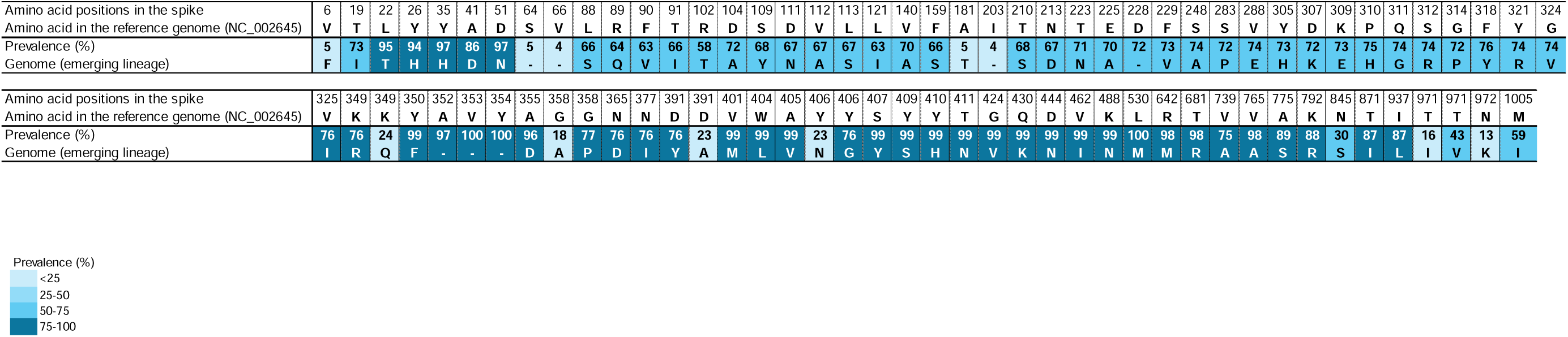
Mutations in the spike protein present in ≥5 genomes compared with reference genome NC_002645 dating backing to 1962.

|  |  |  |  |  |  |  |  |  |  |  |  |  |  |  |  |  |  |  |  |  |  |  |  |  |  |  |  |  |  |  |  |  |  |  |  |  |  |  |  |  |  |  |  |
| --- | --- | --- | --- | --- | --- | --- | --- | --- | --- | --- | --- | --- | --- | --- | --- | --- | --- | --- | --- | --- | --- | --- | --- | --- | --- | --- | --- | --- | --- | --- | --- | --- | --- | --- | --- | --- | --- | --- | --- | --- | --- | --- | --- |
| Amino acid positions in the spike | 6 | 19 | 22 | 26 | 35 | 41 | 51 | 64 | 66 | 88 | 89 | 90 | 91 | 102 | 104 | 109 | 111 | 112 | 113 | 121 | 140 | 159 | 181 | 203 | 210 | 213 | 223 | 225 | 228 | 229 | 248 | 283 | 288 | 305 | 307 | 309 | 310 | 311 | 312 | 314 | 318 | 321 | 324 |
| Amino acid in the reference genome (NC_002645) | V | T | L | Y | Y | A | D | S | V | L | R | F | T | R | D | S | D | V | L | L | V | F | A | I | T | N | T | E | D | F | S | S | V | Y | D | K | P | Q | S | G | F | Y | G |
| Prevalence (%) | 5 | 73 | 95 | 94 | 97 | 86 | 97 | 5 | 4 | 66 | 64 | 63 | 66 | 58 | 72 | 68 | 67 | 67 | 67 | 63 | 70 | 66 | 5 | 4 | 68 | 67 | 71 | 70 | 72 | 73 | 74 | 72 | 74 | 73 | 72 | 73 | 75 | 74 | 74 | 72 | 76 | 74 | 74 |
| Genome (emerging lineage) | F | I | T | H | H | D | N | - | - | S | Q | V | I | T | A | Y | N | A | S | I | A | S | T | - | S | D | N | A | - | V | A | P | E | H | K | E | H | G | R | P | Y | R | V |

|  |  |  |  |  |  |  |  |  |  |  |  |  |  |  |  |  |  |  |  |  |  |  |  |  |  |  |  |  |  |  |  |  |  |  |  |  |  |  |  |  |  |  |
| --- | --- | --- | --- | --- | --- | --- | --- | --- | --- | --- | --- | --- | --- | --- | --- | --- | --- | --- | --- | --- | --- | --- | --- | --- | --- | --- | --- | --- | --- | --- | --- | --- | --- | --- | --- | --- | --- | --- | --- | --- | --- | --- |
| Amino acid positions in the spike | 325 | 349 | 349 | 350 | 352 | 353 | 354 | 355 | 358 | 358 | 365 | 377 | 391 | 391 | 401 | 404 | 405 | 406 | 406 | 407 | 409 | 410 | 411 | 424 | 430 | 444 | 462 | 488 | 530 | 642 | 681 | 739 | 765 | 775 | 792 | 845 | 871 | 937 | 971 | 971 | 972 | 1005 |
| Amino acid in the reference genome (NC_002645) | V | K | K | Y | A | V | Y | A | G | G | N | N | D | D | V | W | A | Y | Y | S | Y | Y | T | G | Q | D | V | K | L | R | T | V | V | A | K | N | T | I | T | T | N | M |
| Prevalence (%) | 76 | 76 | 24 | 99 | 97 | 100 | 100 | 96 | 18 | 77 | 76 | 76 | 76 | 23 | 99 | 99 | 99 | 23 | 76 | 99 | 99 | 99 | 99 | 99 | 99 | 99 | 99 | 99 | 100 | 98 | 98 | 75 | 98 | 89 | 88 | 30 | 87 | 87 | 16 | 43 | 13 | 59 |
| Genome (emerging lineage) | I | R | Q | F | - | - | - | D | A | P | D | I | Y | A | M | L | V | N | G | Y | S | H | N | V | K | N | I | N | M | M | R | A | A | S | R | S | I | L | I | V | K | I |

The genes with the greatest diversity, regardless of the mutation type, were S, which encodes the spike protein, with 67 amino acid mutations per 1,000 amino acids, then Nsp2, N, Nsp3, which encodes a large multi-domain non-structural protein and is an essential component of the viral replication/transcription complex, and E, with between 26 and 28 amino acid mutations per 1,000 amino acids. The genes with the lowest diversity were NSP7, which is part of the RNA-dependent RNA polymerase complex, and NSP16, which encodes a 2′-O-methyltransferase. The two sublineages a and b each harbored specific amino acid mutations. Positions 349, 358, 391, 406 and 971 of the spike protein were found to harbor two different amino acid substitutions according to the sublineage. Thus, genomes of sublineage a shared the following mutations: Nsp2:H40R; S:K349Q; S:D391A; S:Y406N; and N:E33D (Figures 4,5). Regarding genomes of sublineage b, they shared the 22 following mutations, which were not carried by genomes of sublineage a: Nsp2:A350V; Nsp3:V423M; Nsp3:K428Q; S:T19I; S:F229V; S:S248A; S:V288E; S:Y305H; S:K309E; S:P310H; S:Q311G; S:S312R; S:F318Y; S:Y321R; S:G324V; S:V325I; S:K349R; S:N365D; S:N377I; S:D391Y; S:Y406G; and S:V739A (Figures 4, 5). Among these mutations, 19 were present in the S gene. Four amino acid mutations located in the S gene that we detected in sublineages a or b genomes were previously reported as newly detected^11^.

**Figure 5.**
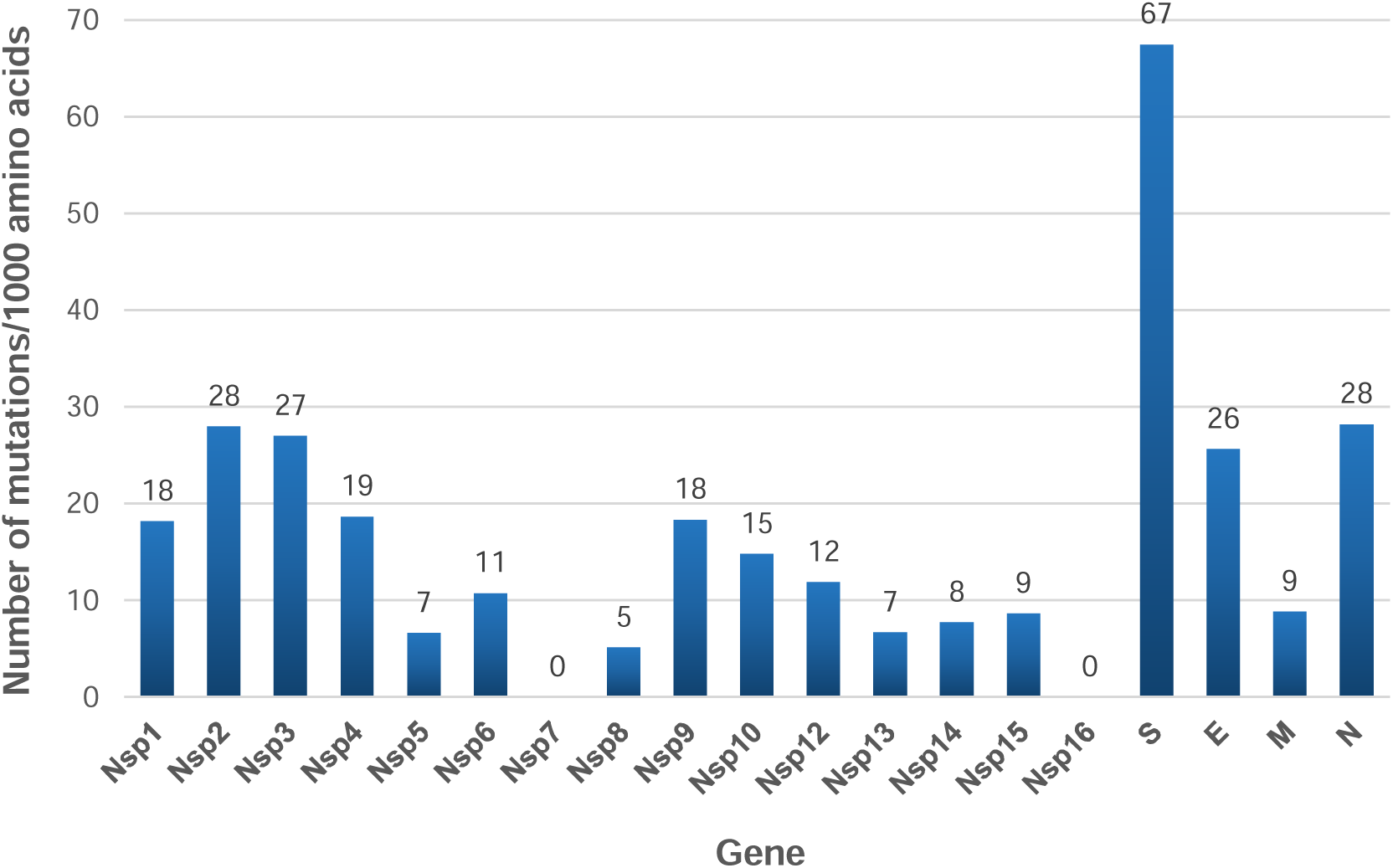
Number of amino acid mutations per 1,000 amino acids Nsp11 was excluded from the analyses as it is only 51 nucleotide-long and overlaps Nsp12. E, envelope; M, Membrane; N, Nucleocapsid; S, Spike.

## DISCUSSION

We report here a multiplex PCR system to amplify whole HCoV-229E genomes before their sequencing by next-generation technologies. The present study allowed obtaining and analyzing 123 new near full-length HCoV-229E genomes, which was a number almost equal to the global number of HCoV-229E genomes at the beginning of Year 2024. In addition, we provided here the first HCoV-229E genomes from patients sampled in France. Another similar multiplex PCR systems for HCoV-229E whole genome amplification were reported in 2024 by Musaeva et al^31^ and McClure et al^30^, in Russia and UK, respectively. The first one^31^ used twenty-nine PCR primer systems, compared to 31 in the present study, and enabled obtaining 39 genomes with a coverage of a full-length genome >70%, from 50 HCoV-229E RNA-positive nasopharyngeal swab samples. The second one^30^ used 29–36 primer pairs for all four endemic human coronaviruses (HCoV-229E, -OC43, -NL63, and −229E), and enabled obtaining 64 HCoV-229E genomes with a coverage of a full-length genome >95%. Overall, as of 01/02/2025, 269 HCoV-229E genomes were available in GenBank, and the 123 genomes obtained here grew the global set to 392.

Most of the HCoV-229E genomes obtained here were retrieved from nasopharyngeal samples collected between 2019 and 2022. This can be explained by the Covid-19 pandemic that led to a massive diagnosis of respiratory virus infections during this period of time. Between July 2020 and June 2021, few HCoV-229E infections were diagnosed in our center, as previously reported, which could be explained at least partly by the implementation of hygiene measures to fight the SARS-CoV-2 pandemic^32^. All the genomes obtained here belong to the emerging lineage previously reported by Ye et al^11^, with 43 full-length genomes, who demonstrated that this lineage circulated between 2016 and 2020 in different countries and suggested that it was the majority lineage circulating in the world. We report here that this lineage, which was tentatively reported as genotype 7 in Musaeva et al’s and McClure et al’s study^30,31^, also circulated in France. Two sublineages 7a and 7b had been reported for this genotype^30,31^, while we also observed in the present study two sublineages “a” and “b” with an overall predominance of sublineage b. Hence, our findings further support that the emerging lineage initially reported in China, then in Japan, Haiti, the United States, Russia, and UK likely became the predominant lineage worldwide. In addition, our findings together with previous data,^11,30,31^ indicate that this emerging lineage are evolving with new mutations whose occurrence may depend on the time and geographical area of HCoV-229E circulation. Sublineages in this emerging lineage should be more precisely delineated in further studies with additional HCoV-229E genomes.

A high amino acid diversity was observed here in the spike and non-structural proteins, which is consistent with previous findings in coronaviruses^11,30,31,33–35^. The HCoV-229E receptor binding domain (RBD) contains three loops, named 1, 2 and 3, which are involved in the binding of the virus to the host aminopeptidase N cellular receptor and are located at amino acid positions 308-325, 352-359 and 404-408, respectively^34^. Some amino acid mutations in these regions were observed here as in three previous studies^11,30,31,35^. These notably involved four amino acid positions. For three of them, two different mutations were observed here (K349R or Q; G358P or A; Y406G or N). For position 391, two types of substitutions were observed here that are signatures of either sublineage a (D391A) or sublineage b (D391Y). Besides, a tryptophan at position 404 of the spike RBD was reported to be very important for loop 3 binding to the cellular receptor^34^. Notwithstanding, tryptophan was replaced here by a leucine in all genomes, indicating that this latter does not preclude viral infection. Mutations Q430K, D444N and K488N also encountered in the present study were already reported and would result in an N-glycosylation site at position 488^35^. Besides, apolar bonds were predicted that involve certain amino acids in the RBD, notably at position 318^34^. A mutation at this position was reported to be associated with an ≈13-fold reduction in the affinity with the cellular receptor^34^. Here, a mutation (F318Y) at this position was found in 76% of the genomes. This same mutation was also reported previously in HCoV-229E genotypes 3, 4, 5, 6 and in the emerging lineage as well^11^. Taken together, these data highlight the broad diversity of spike amino acid patterns and may be useful for the interpretation of structural analyses performed previously and in future studies including to investigate the putative impact of these different mutations on the interaction between HCoV-229E and the host receptor^36–38^.

Overall, the present study and two other recent studies^30,31^enrich the set of HCoV-229E genomes available worldwide, with 123 and 103 genomes respectively. Nonetheless, genomic data remain scarce and they cover a limited number of countries with still limited data sets, therfore not necessarily reflecting the circulation of this virus at the global scale. Further studies are therefore needed to gain a more global view of the evolutionary dynamics of HCoV-229E, which will clarify the specificities of this virus and may contribute to a more general understanding of the evolution of human coronaviruses.

## Acknowledgments

We are very grateful to the technical team of the next-generation sequencing platform of IHU Méditerranée Infection.

## Data Availability

HCoV-229E genomes analysed here have been submitted to the GenBank sequence database^20^ (https://www.ncbi.nlm.nih.gov/genbank/) (GenBank Submission ID 2929308).

## Author contributions

Conceived and designed the experiments: BLS, PC. Contributed materials, analysis tools: All authors. Analyzed the data: HH, JP, EB, PC. Writing—original draft preparation: HH, JP, PC. Writing—review and editing: All authors. All authors have read and agreed to the published version of the manuscript.

## Ethics

The present study has been registered on the Health Data Access Portal of Marseille public and university hospitals (Assistance Publique-Hôpitaux de Marseille (AP-HM)) with No. PADS24-190 and was approved by the Ethics and Scientific Committee of AP-HM.

